# Visual familiarity learning at multiple timescales in the primate inferotemporal cortex

**DOI:** 10.1101/2024.01.05.574412

**Authors:** Krithika Mohan, Ulises Pereira-Obilinovic, Stanislav Srednyak, Yali Amit, Nicolas Brunel, David Freedman

## Abstract

Humans and other primates can rapidly detect familiar objects and distinguish them from never-before-seen novel objects. We have an astonishing capacity to remember the details of visual scenes even after a single, fleeting experience. This ability is thought to rely in part on experience-dependent changes in the primate inferotemporal cortex (IT). Single neurons in IT encode visual familiarity by discriminating between novel and familiar stimuli, with stronger neural activity on average for novel images. However, key open questions are to understand how neural encoding in IT changes as images progress from novel to highly familiar, and what learning rules and computations can account for learning-dependent changes in IT activity. Here, we investigate the timescales over which novel stimuli become familiar by recording in IT as initially novel images become increasingly familiar both within and across days. We identified salient and persistent memory-related signals in IT that spanned multiple timescales of minutes, hours, and days. Average neural activity progressively decreased with familiarity as firing rates were strongest for novel, weaker for intermediately familiar, and weakest for highly familiar images. Neural signatures of familiarity learning were slow to develop as response reductions to initially-novel images emerged gradually over multiple days (or hundreds of views) of visual experience. In addition to slow changes that emerged across sessions, neural responses to novel images showed rapid decreases with familiarity within single sessions. To gain insight into the mechanisms underlying changes of visual responses with familiarity, we use computational modeling to investigate which plasticity rules are consistent with these changes. Learning rules inferred from the neural data revealed a strong diversity with many neurons following a ‘negative’ plasticity rule as they exhibited synaptic depression over the course of learning across multiple days. A recurrent network model with two plasticity time constants – a slow time constant for long timescales and a fast time constant for short timescales – captured key dynamic features accompanying the transition from novel to familiar, including a gradual decrease in firing rates over multiple sessions, and a rapid decrease in firing rates within single sessions. Our findings suggest that distinct and complementary plasticity rules operating at different timescales may underlie the inferotemporal code for visual familiarity.

## 1 Introduction

Primates experience objects within a rich spatial and temporal context, perceiving at once whether a fruit or a face is familiar or never seen before. Remarkably, we have the capacity to learn and remember an enormous number of familiar visual objects (Standing, 1973). This includes items that were viewed very recently (minutes or hours ago), as well as objects seen only days, months, or even years earlier (Bahrick et al., 1975). This perceptual and mnemonic ability is crucial for our ability to recognize and recall familiar patterns, items, and individuals, and is thus a fundamental ingredient for creating a meaningful internal model of the world.

Visual familiarity is encoded in the activity of single neurons and populations in the primate inferotemporal cortex (IT), a key node in the ventral visual stream that is critical for object recognition (Li et al., 1993; Miyashita, 1993; Tanaka, 1996). Neural activity in IT is correlated with visual familiarity as single neurons often encode novel, never-before-seen images with higher average firing rates than familiar images that have been seen thousands of times (Fahy et al., 1993; Sobotka and Ringo, 1993; Xiang and Brown, 1998). Indeed, IT populations show reduced firing rates even after a single exposure to novel images in visual memory tasks (Meyer and Rust, 2018). While familiar images evoke lower firing rates on average than novel images, neural selectivity to familiar images is often greater than for novel images despite lower average firing rates (Freedman et al., 2006; Woloszyn and Sheinberg, 2012a). It has also been reported that familiarity with an image can produce highly specific increases in firing rates among neurons which have a strong preference for that preferred image (Woloszyn and Sheinberg, 2012a). This suggests a hypothesis that those neurons which are initially strongly driven by a stimulus may show enhanced firing rates as that stimulus becomes familiar, while other neurons (which are not specifically tuned for that image) show depression with familiarity (Hebb, 1949). These neural signatures of familiarity can be quantitatively captured by unsupervised Hebbian models of learning, provided the underlying learning rule is dominated by synaptic depression, which accounts for the progressive decrease in average visual responses with familiarity (Amit, 1995; Lim et al., 2015).

Such experience-dependent changes in IT circuits have been examined by comparing only two classes of stimuli, entirely novel and highly familiar, and have therefore only provided an asymptotic view of learning, with no information about the timescales of this learning process. But how and when IT neurons show changes in neural encoding as novel stimuli become familiar has remained a mystery. What happens to the neural code of initially novel stimuli as they become familiar? What plasticity rules govern the transition from novel to familiar?

Here we investigate the neural dynamics of visual learning at multiple timescales – starting from short within-session timescales of single image presentations up to long timescales of thousands of image presentations. We recorded from IT cortex in monkeys during a task that involved successive presentations of large numbers of initially novel stimuli that were in turn presented many times on successive days. This task enabled us to examine changes in neural encoding as initially novel stimuli became increasingly familiar over the course of a recording session, and across multiple daily recording sessions.

IT circuits carry a robust signal of graded visual familiarity with strongest responses to novel images, followed by a progressive decrease in average responses to intermediately familiar and highly familiar images. Neural correlates of familiarity developed gradually across many image presentations over multiple days, with a modest impact of short-term within-session familiarity compared to large impacts of long-term familiarity. Plasticity rules fitted to data exhibit a large heterogeneity, with a dependence on the post-synaptic firing rate that can be either purely negative, positive, Hebbian or anti-Hebbian. Finally, we show that data can be captured by a computational model with two different time constants of synaptic plasticity, one on the order of minutes-to-hours, the other on the order of days-to-weeks. Our combined experimental and theoretical approach unveils the multiscale dynamics of learning in cortical circuits and provides key mechanistic insights into the plasticity rules operating at both short and long timescales.

## 2 Results

### 2.1 Dimming detection task

To investigate the timescales of visual learning in IT, we first familiarized two monkeys with images from a large library (∼500 images) comprising natural and man-made, objects and scenes. During familiarization, monkeys were trained to perform an image luminance-change or “dimming” detection task in which they detected and reported a decrease in image luminance in a series of rapidly presented images (Figure 1). The purpose of the dimming detection task was to ensure that the animals were actively attending to each image, and to encourage them to be similarly task-engaged across the session. The images were presented at central fixation for 400 ms each with a 150 ms inter-stimulus interval. On each trial, the number of images presented before the dimmed image ranged from one to five. On half of the trials, the luminance of one of the images decreased (i.e., the image dimmed) and the monkeys were rewarded for releasing a manual touch bar when they detected dimming. On the remaining half of the trials, five images were always presented and none of the images presented in the sequence dimmed, and the monkeys were rewarded for withholding a response. Both monkeys successfully performed the dimming detection task with a mean accuracy of 99% and a mean reaction time of 326 ms on dimming trials (Monkey B: accuracy 99.26%, reaction time 313 ms; Monkey S: accuracy 98.02%, reaction time 350 ms).

**Figure 1:**
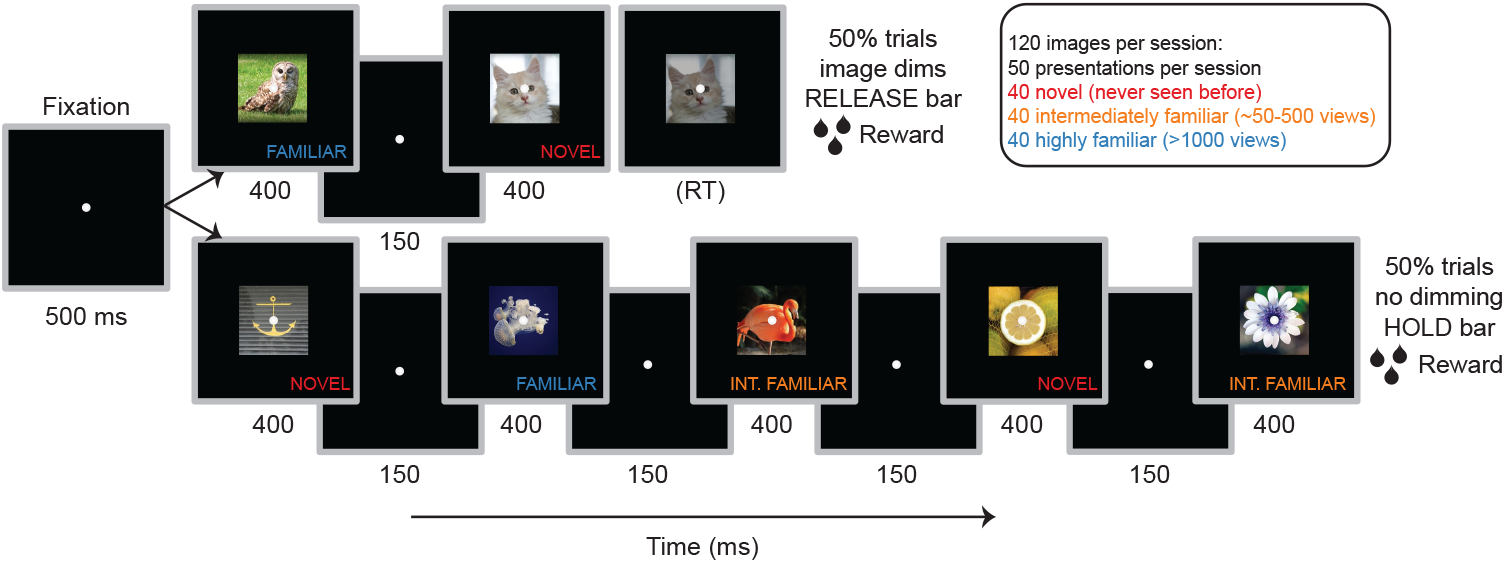
RSVP dimming detection task. Monkeys detected a decrease in the luminance of an image (i.e. dimming) and reported dimming through the release of a manual touch-bar. Each trial started with a 500 ms fixation period, followed by image presentation for 400 ms with an inter-stimulus interval of 150 ms. On half of the trials, up to five images were shown and dimming occurred on any of the five images and monkeys were rewarded for releasing the touch-bar to report image dimming (top: example trial depicts dimming on the second image). On the remaining half of the trials, five images were always presented and no dimming occurred and monkeys were rewarded for holding the touch-bar through the trial (bottom panel). The images varied in their levels of familiarity to the monkeys. They were either novel (never seen before), intermediately familiar (∼50-500 views), or highly familiar (*>*1000 views) to the monkeys and were randomly interleaved within and across trials.

To examine how neural codes of familiarity evolved over long and short timescales, we tested novel images with two groups of familiar images at different levels of familiarity – highly familiar and intermediately familiar images. The highly familiar images were randomly sampled from a library of familiarized images and were previously viewed at least 1000 times over 3-5 months during long-term familiarization prior to the start of recordings. The intermediately familiar images were repeated over a week of consecutive recording sessions and spanned a range of 50-400 views (this corresponded to approximately 50 views each day for 8 days). As the intermediately familiar images were presented over multiple days during recordings, they enabled us to ask how familiarity-related changes emerged over long timescales of days to weeks. Finally, the novel images consisted of unique images that were never seen before each recording session, were viewed approximately 50 times within a single session and replaced in subsequent recording sessions. Since the novel images were presented multiple times within a session, they allowed us to ask how familiarity representations changed over short timescales of minutes to hours, i.e., within a single session. In each recording session, 120 unique (40 novel, 40 intermediately familiar and 40 highly familiar) images were presented about 50 times in the dimming detection task. Thus, the images varied in their familiarity to the monkeys as follows: (i) novel (never seen before that recording), (ii) intermediately familiar (viewed 50-500 times over one week during consecutive recording sessions), and (iii) highly familiar (viewed at least 1000 times over 3-5 months before recording).

### 2.2 Graded image familiarity is represented in single IT neurons

We recorded from IT neurons using 24-channel linear probes while two monkeys performed the dimming detection task with novel, intermediately familiar, and highly familiar images (Monkey B: N=306 single units, Monkey S: N=233 single units). To determine whether IT neurons encode image familiarity at multiple timescales, we compared responses of well-isolated, visually-selective neurons to the three image categories. Out of 539 recorded neurons, 86% of neurons were visually selective (Monkey B: 82%, Monkey S: 92%, *p <* 0.05, Linear model, see Methods). Many of these visually selective IT neurons encoded image familiarity in a graded manner with most neurons responding with highest firing rates to novel, lower firing rates to intermediately familiar, and lowest firing rates to familiar images (Figure 2a, 2b). Consistent with this, average population responses in IT revealed a similar progression, with higher firing rates to novel, followed by intermediately familiar, and then highly familiar images (Figure 3a). We found that 40% of neurons were significantly modulated by whether an image was familiar or not, independent of the visual selectivity for individual images (Monkey B: 37%, Monkey S: 43%; Linear model *p <* 0.05). While most neurons showed strongest firing rates to novel images, a smaller number (16.5%) of familiarity-selective neurons showed the opposite trend responding with higher firing rates to either highly familiar or intermediately familiar images, and lowest to novel images (Figure 2c) (Tukey’s HSD criterion, p*<*0.05). Thus, average population activity in IT reflects graded image familiarity through a continued decrease in neural activity with increasing levels of familiarity at multiple timescales of hours (novel), weeks (intermediately familiar), and months (highly familiar).

**Figure 2:**
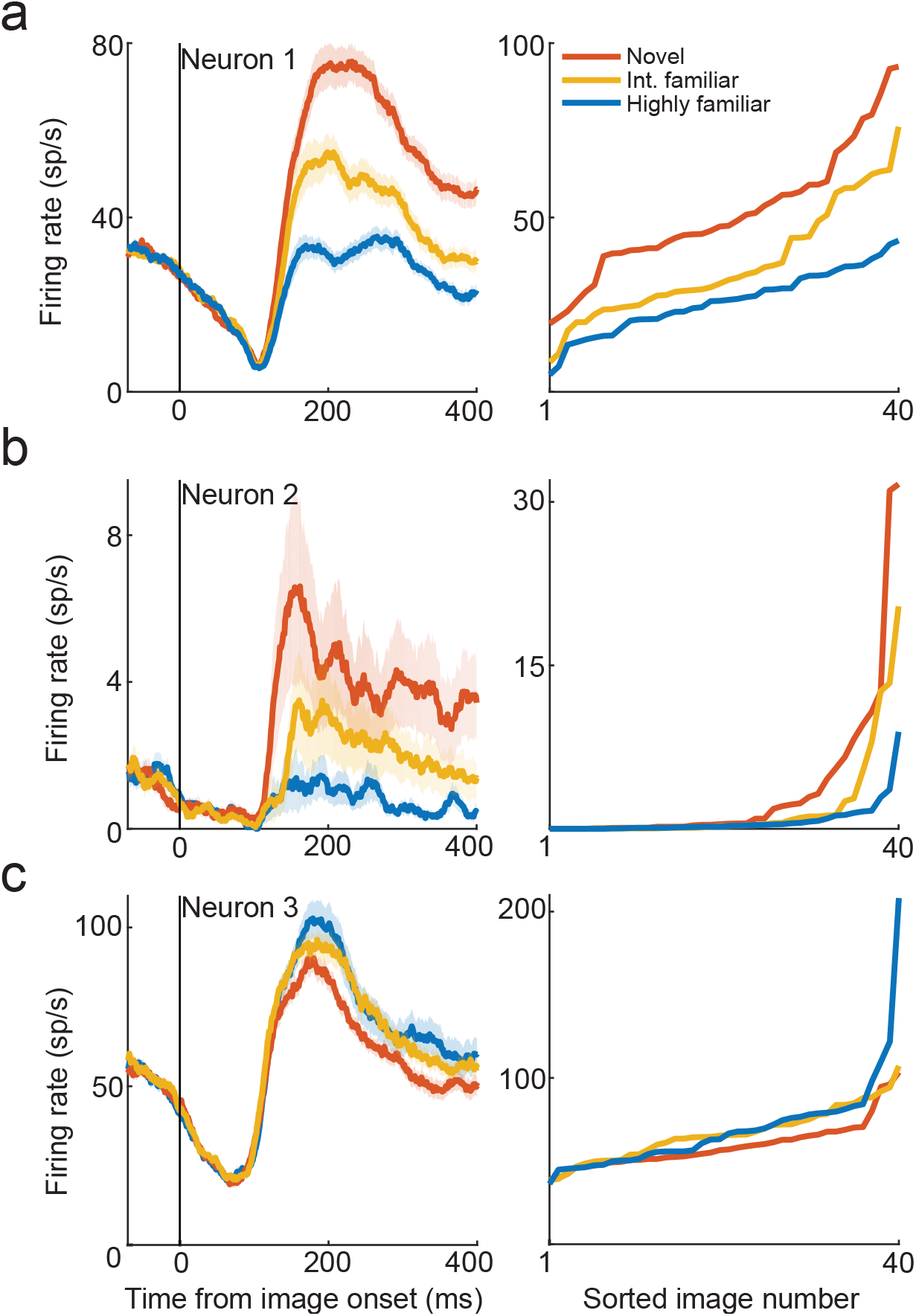
Single IT neurons encode graded levels of image familiarity. (a) Left: Peri-stimulus time histogram (PSTH) of a visually selective IT neuron to novel (seen ∼10s of times), intermediately familiar (seen ∼100s of times), and highly familiar images (seen ∼1000s of times). Average firing rates are strongest for novel, followed by intermediately familiar, and weakest for highly familiar images. Colors represent different levels of familiarity. Vertical line at 0 indicates image onset. Spike trains were convolved with a 20-ms causal boxcar kernel. Right: Rank-ordered curves of the firing rates of the corresponding single neuron on the left to novel, intermediately familiar, and highly familiar images. Firing rates are averaged over the image presentation period (80-480 ms from image onset) (b) PSTH of another visually selective, sparse IT neuron, with highly selective responses to a small set of images. (c) PSTH of another visually selective IT neuron, with strongest firing rates for highly familiar, followed by intermediately familiar, and weakest for novel images. Both (b) and (c) follow plotting conventions as in (a).

**Figure 3:**
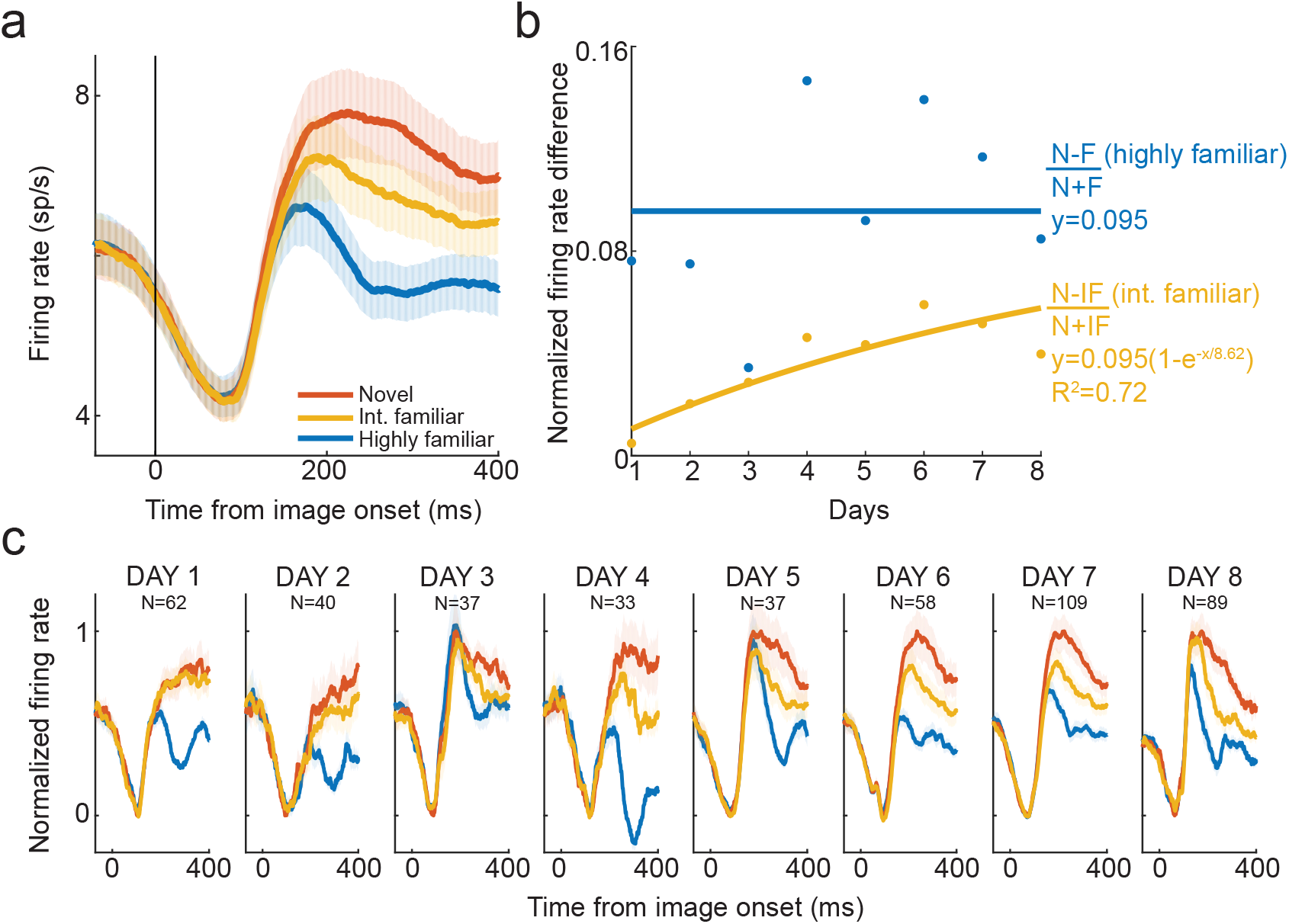
Long-term image familiarity emerges slowly over multiple days in IT neural populations. (a) Peri-stimulus time histogram (PSTH) of visually selective IT neurons to novel (seen ∼10s of times), intermediately familiar (seen ∼100s of times), and highly familiar images (seen ∼1000s of times). Average firing rates are strongest for novel, followed by intermediately familiar, and weakest for highly familiar images. (b) Normalized firing rate difference between responses to novel and intermediately familiar images (N-IF/N+IF in orange) as a function of number of days of viewing intermediately familiar images. The blue curve represents the normalized firing rate difference between responses to novel and highly familiar images (N-F/N+F) as a function of number of days of viewing highly familiar images. Positive values indicate that neural responses to novel images are greater than responses to both intermediately familiar and highly familiar images. Over multiple days of viewing, the normalized firing rate difference between novel and intermediately familiar images increases, following an exponential function with an asymptote that reaches the normalized difference in firing rates between novel and highly familiar images. (c) Normalized PSTHs to novel, intermediately familiar and highly familiar images over multiple days of repeated image presentations. The difference between novel and intermediately familiar images increases over days, reaching statistical significance first on day four and remaining significant until day eight. For each day, the PSTH represents the firing rates pooled over all images in a given category and across all neurons recorded on that day (depicted above the PSTHs for each day). Firing rates on each day were normalized by subtracting the minimum firing rate and dividing by the maximum minus minimum firing rate, with both minimum and maximum firing rates measured during the image presentation period (80 to 480 ms from image onset). In both (a) and (c), spike trains were convolved with a 20-ms causal boxcar kernel, and shaded error bars indicate SEM.

### 2.3 Long-term image familiarity emerges slowly over multiple days in IT neural populations

We next asked how neural representations of image familiarity changed with long-term visual experience over several days. To characterize how neural activity evolved over multiple days, we quantified how responses to intermediately familiar images changed as they were repeatedly presented over eight consecutive days of recording. The intermediately familiar images in our image set spanned a wide range of views, from about 50 views on the first day of recording to 400 views on the eighth day of recording (with about 50 views per day). Since we recorded from non-overlapping IT sub-populations in every session, we compared neural responses to intermediately familiar images in relation to novel images presented during the same recording session.

Neural responses to intermediately familiar images gradually decreased over several days and consistently remained lower than responses to novel images and higher than responses to highly familiar images (Figure 3c). The difference between responses to novel and intermediately familiar images increased over successive days of repeated presentations, suggesting that intermediately familiar images gradually became familiar. We quantified the rate at which neural activity to intermediately familiar images decreased over days by calculating the normalized difference in firing rates for each session as (N-IF)/(N+IF) where N (or IF) is the mean firing rate across all neurons to novel (or intermediately familiar) images (Figure 3b, orange line). To infer when, i.e., on what day this firing rate difference becomes significant, we fit a generalized linear model to neural firing rates as a function of day, image category (N and IF), and neuron number. We asked whether firing rates to novel and intermediately familiar images significantly differed from each other depending on how many days they had been viewed. The model first achieved a significant interaction between day and image category on day four (or 200 views) of repeated presentations, and remained significant on all subsequent days until day eight in both monkeys (*p* = 0.01 for day 4, *p <* 0.05 for days 5-8, interaction between day x image-category, GLM, see Methods and Table 1). Moreover, this normalized firing rate difference increased smoothly over multiple days, showing that long-term familiarity of the order of 100s of views is accompanied by a slow, progressive reduction in neural activity.

**Table 1:** Number of neurons whose fitted parameters are significantly outside a 95% or 99% confidence interval, as determined by randomly shuffling visual responses. Visual responses computed from the early response (75-200ms)

| Type | Criterion | Number of neurons | Expected number | Probability |
| --- | --- | --- | --- | --- |
| ‘Negative’ | $a_c < a_{c05}$ | 63 | 23.25 | $< 10^{-6}$ |
| | $a_c < a_{c01}$ | 22 | 4.65 | $< 10^{-6}$ |
| ‘Positive’ | $a_c > a_{c95}$ | 34 | 23.25 | 0.02 |
| | $a_c > a_{c99}$ | 10 | 4.65 | 0.02 |
| ‘Anti-Hebbian’ | $b_l < b_{l05}$ | 42 | 23.25 | 0.0002 |
| | $b_l < b_{l01}$ | 11 | 4.65 | 0.008 |
| | $b_s < b_{s05}$ | 37 | 23.25 | 0.004 |
| | $b_s < b_{s01}$ | 8 | 4.65 | 0.1 |
| ‘Hebbian’ | $b_l > b_{l95}$ | 33 | 23.25 | 0.03 |
| | $b_l > b_{l99}$ | 9 | 4.65 | 0.05 |
| | $b_s > b_{s95}$ | 29 | 23.25 | 0.13 |
| | $b_s > b_{s99}$ | 8 | 4.65 | 0.1 |

We wondered whether this response reduction was a signature of familiarity-related learning, was it specific to images that were intermediately familiar or was it a form of generalized visual habituation towards all image types independent of familiarity? To test this, we assessed whether neural responses to highly familiar images, that were previously seen 1000s of times, also decreased with repeated presentations over several days. We hypothesized that highly familiar images – that were experienced 1-2 orders of magnitude more than the other images – would not elicit a continued decrease over multiple days of repeated viewings. Since we included highly familiar images in every recording session along with novel and intermediately familiar images, we similarly quantified the change in firing rates to highly familiar images relative to novel images from the same session. As described above, we calculated the normalized difference in firing rates for each session as (N-F)/(N+F) where F is the mean firing rate across all neurons to highly familiar images in each session. In contrast to the responses to intermediately familiar images, the normalized firing rate difference between novel and highly familiar images remained constant over days and was well fit by a constant function (Figure 3b, blue line, linear model with zero slope, intercept=0.095), indicating that firing rates tend to saturate once images have become highly familiar. Our results suggest that the observed response suppression for intermediately familiar images (but not highly familiar images) reflects an ongoing process of familiarity-related learning that reaches an asymptote for highly familiar, learned images.

Finally, we sought to understand the timescales over which intermediately familiar images become highly familiar. Specifically, we set out to quantify how long it takes for IT neural activity to reach an asymptote i.e., at what point does any further increase in the number of views no longer cause a familiarity-related decrease in firing rates? To infer the timescales at which neural responses saturate, we evaluated the rate of change of the firing rate difference to novel and intermediately familiar images over days. The index (N-IF/N+IF) initially increased over the first few days and appeared to level off to an asymptote (Figure 3b). The rate of change of this index was modeled using an exponential decay function with an asymptote. The asymptote for this model was set to match the constant value obtained for highly familiar images, specifically, the mean of (N-F/N+F) over days. The exponential model yielded a time constant of nine days, revealing that it would take more than a week of ∼50 repeated viewings of initially novel images each day for IT firing rates to saturate and not decrease further. These results highlight two findings: first, progressive response reduction is a neural correlate of familiarity-related learning and second, neural representations of long-term image familiarity develop slowly over a week of repeated presentations of initially novel images.

### 2.4 Short-term image familiarity emerges rapidly over multiple hours in IT neural populations

Our results so far reveal that IT neurons encode degrees of familiarity, and that IT activity on average decreases slowly as images become increasingly familiar over multiple days. However, it remains unclear how familiarity-related responses change with repeated image presentations within the time scale of minutes to hours in a single recording session. To examine this, we focus our analyses on visually selective neurons within single sessions that were present from the start of the recording session and remained well-isolated for up to at least 25 presentations of each image. We note that as repetitions increased beyond 25, the number of neurons that remained well-isolated from the start of recording falls off markedly due to drift, and thus we selected 25 presentations to include a reasonable number of neurons (this corresponded to about 1.5 hours into the recording session) (Monkey B: N=106 single units, Monkey S: N=78 single units, Figure S1). Within single sessions, average firing rates to novel images decreased over repetitions with a pronounced drop in the first few presentations, whereas average firing rates to highly familiar images remained near-constant with repeated presentations (Figure 4a, 4b). The decrease in firing rates for both novel and familiar images was captured by linear models with negative slopes (Novel: slope = -0.04, intercept = 3.85, p=10^*−*7^; Familiar: slope = -0.009, intercept = 2, p=0.02). We tested whether within-session firing rates decreased at different rates for novel and familiar stimuli and found that firing rates within a session decreased more rapidly for novel than for familiar images (ANCOVA for novel versus familiar, p=0.0001). As the session progressed, the difference in firing rates between novel and familiar images gradually decreased, but it never reached zero. This observation implies that representations of familiarity in IT do not emerge abruptly within one to two hours in a single session. Rather, IT activity tracks image familiarity over repetitions that span several orders of magnitude ranging from tens to hundreds to thousands of image presentations.

**Figure 4:**
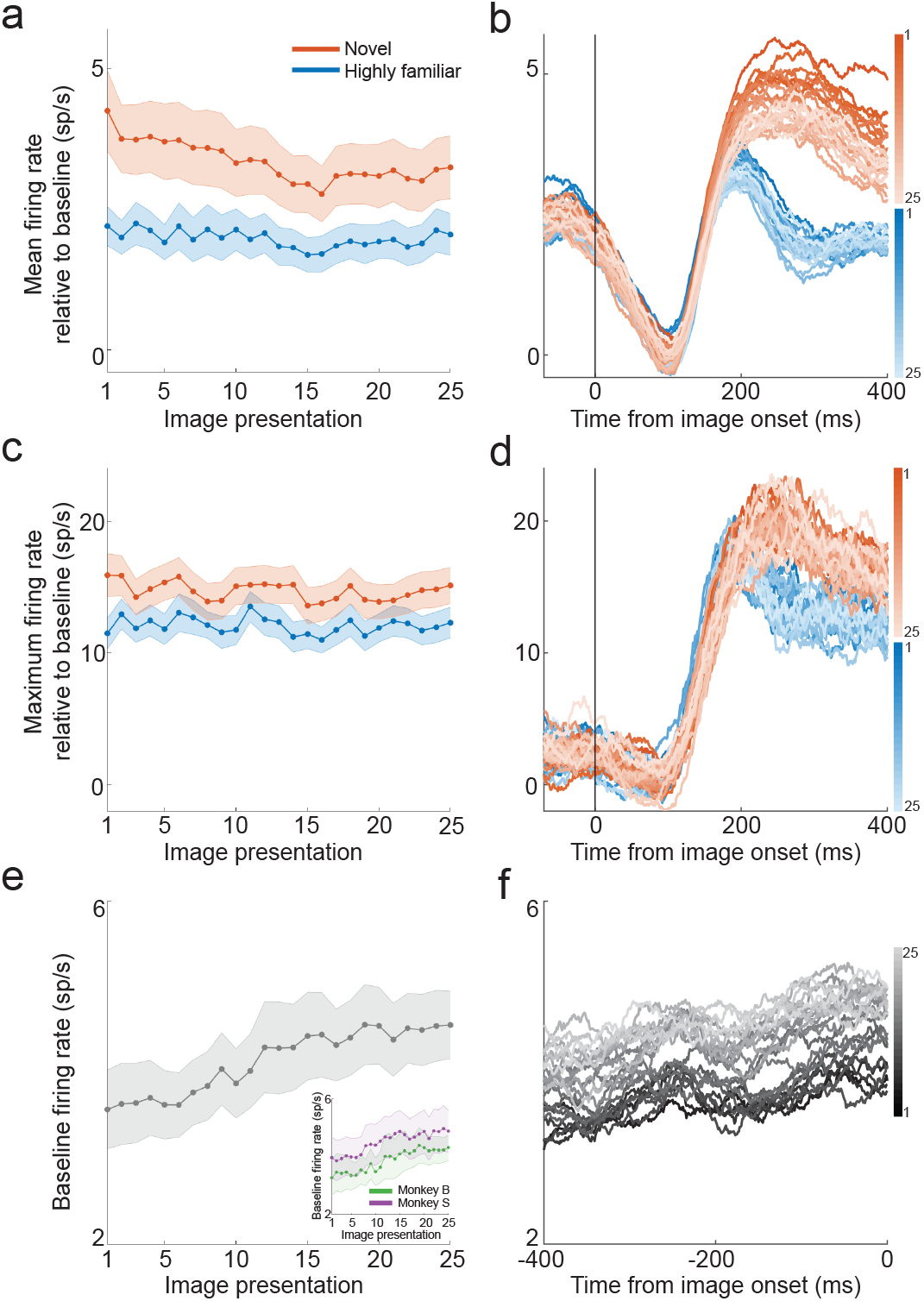
Short-term image familiarity emerges rapidly within a session in IT neural populations. (a) Mean neural activity of visually selective IT neurons to all novel and all highly familiar images over repeated image presentations within a session. (b) Time course of mean neural activity of visually selective IT neurons to novel and highly familiar stimuli. Average firing rates decreased significantly more to novel than to highly familiar stimuli over repetitions within a session. (c) Maximum neural activity of visually selective IT neurons to the best novel and the best familiar image over repeated image presentations within a session. (d) Time course of maximum neural activity of visually selective IT neurons to the best novel and best familiar stimulus. Maximum firing rates remained stable to both novel and highly familiar stimuli over repetitions within a session. (e) Pre-stimulus baseline activity of visually selective IT neurons over repeated image presentations within a session. Firing rates are averaged over the fixation period (400 ms preceding image onset). Inset: Baseline firing rates over repeated presentations within a session for both monkeys. (f) Time course of baseline firing rates during the fixation period. Baseline firing rates increase with repeated presentations within a session. In panels (a) and (c), firing rates were pooled over all images in a given category and across all neurons. Mean firing rates were computed by averaging neural activity over the image presentation period (80-480 ms from image onset) and subtracting baseline firing rates averaged over the pre-stimulus period (400 ms preceding the onset of the first image in that trial). In panels (b), (d) and (f), firing rates were smoothed using a 50-ms causal boxcar kernel, colored by familiarity and shaded by repetition within a session.

In the previous analysis, we examined how familiarity affects neural activity in IT by averaging across all novel and familiar stimuli. However, neurons in IT exhibit sparse selectivity, often responding maximally to a small number of images while being unresponsive to many others. Moreover, neurons that are most responsive (to a particular stimulus) are likely to have a more significant impact on readout by downstream neurons, highlighting their role in models of readout of object representations. How does familiarity within a session affect neural responses to neurons’ most preferred novel and familiar stimuli? One possibility is that IT responses to the most preferred novel stimuli (but not familiar stimuli) may increase with repeated presentations in a session resulting in greater selectivity, and this rapid increase in selectivity could underlie familiarity learning at short timescales. Alternatively, neural responses to the ‘best’ novel stimuli might remain relatively unchanged within a session, indicating that visual experience has minimal impact on neural selectivity over within-session timescales. To investigate these possibilities, we analyzed neural responses for the best or most preferred stimuli for each neuron. Specifically, for each neuron we computed firing rates produced by two images: the most preferred novel image that produced the highest mean firing rate, and the most preferred familiar image that produced the highest mean firing rate. We found that the maximum firing rates to the best novel and the best familiar images remained near-constant over repetitions within a session (linear fit for best novel: slope = -0.04, intercept=15.19, p=0.05; linear fit for best familiar: slope = -0.01, intercept=12.3, p=0.3), suggesting that short-term familiarity had little to no impact on the visual selectivity of IT neurons (Figure 4c, 4d). However, firing rates to the best novel stimuli consistently remained greater than the best familiar stimuli throughout the session, suggesting that longer time scale familiarity learning produces reliable differences in IT firing rates for novel versus highly familiar stimuli which remain distinct over the one to two hour within-session time scale.

Our findings hint at the idea that visual familiarity learning in IT cortex is mediated by multiple mechanisms which operate over different timescales. At short timescales (i.e., minutes to hours), neural activity to novel stimuli decreased in the first few repetitions after which it remained near constant, yet distinct from familiar stimuli. On the other hand, at longer timescales (i.e., days to weeks), activity steadily decreased with repeated presentations and long-term familiarity emerged over the course of a week. How is it that these distinct patterns of firing rate changes within single sessions gives rise to learning and plasticity that manifests slowly across multiple days? One explanation is that, over short timescales, learning - specifically, continued visual experience with novel stimuli - could increase excitability of neural ensembles over a time period that coincides with memory consolidation, thus facilitating synaptic plasticity over long timescales (Chen et al., 2020). Enhanced excitability has been observed in the rodent hippocampus after learning various tasks including fear conditioning and olfactory discrimination (Cai et al., 2016; Zelcer et al., 2006). A recent study also found that spontaneous cortical activity (but not visually evoked activity) in mouse V1 increased after visual habituation and that optogenetically boosting baseline activity accelerated habituation, suggesting that spontaneous activity could play a direct role in visual learning (Miller et al., 2022). To investigate this possibility, we used baseline firing rates (400 ms prior to image onset) as a proxy for neural excitability and measured how they changed within single sessions. Interestingly, baseline firing rates showed a prominent increase with repeated presentations in single sessions, an effect that was robust in both monkeys (Figure 4e, 4f). Notably, this increase in neural activity occurred only in the pre-stimulus baseline activity, whereas visually evoked activity to novel or familiar images showed a decreasing trend or remained constant. Together, these observations suggest a role for intrinsic plasticity in learning by which an increase in neural excitability over timescales of hours might temporally overlap with synaptic consolidation and contribute towards the emergence of familiarity memory over timescales of days.

### 2.5 Inference of plasticity rules from data

To gain a deeper understanding of the circuit mechanisms underlying familiarity learning, we next asked what plasticity rules could lead to the observed changes in visual response statistics with familiarity. One possibility is that these changes occur due to changes in synaptic connectivity in IT cortex. This hypothesis was strengthened by Lim et al. (2015) who showed that changes in distributions of visual responses with familiarity can be explained by an unsupervised synaptic plasticity rule that changes synaptic strengths between neurons as a function of the firing rates of pre and post-synaptic neurons, Δ*J*_*ij*_ = *f* (*r*_*i*_)*g*(*r*_*j*_) where *r*_*i*_ and *r*_*j*_ are the firing rates of the post and the pre-synaptic neurons, respectively. They developed a procedure to infer the dependence of the plasticity rule on the post-synaptic firing rate, *f* (*r*_*i*_) from distributions of visual responses to novel and familiar stimuli. This procedure is explained in detail in Methods and is illustrated in Fig. 5 for one example neuron.

**Figure 5:**
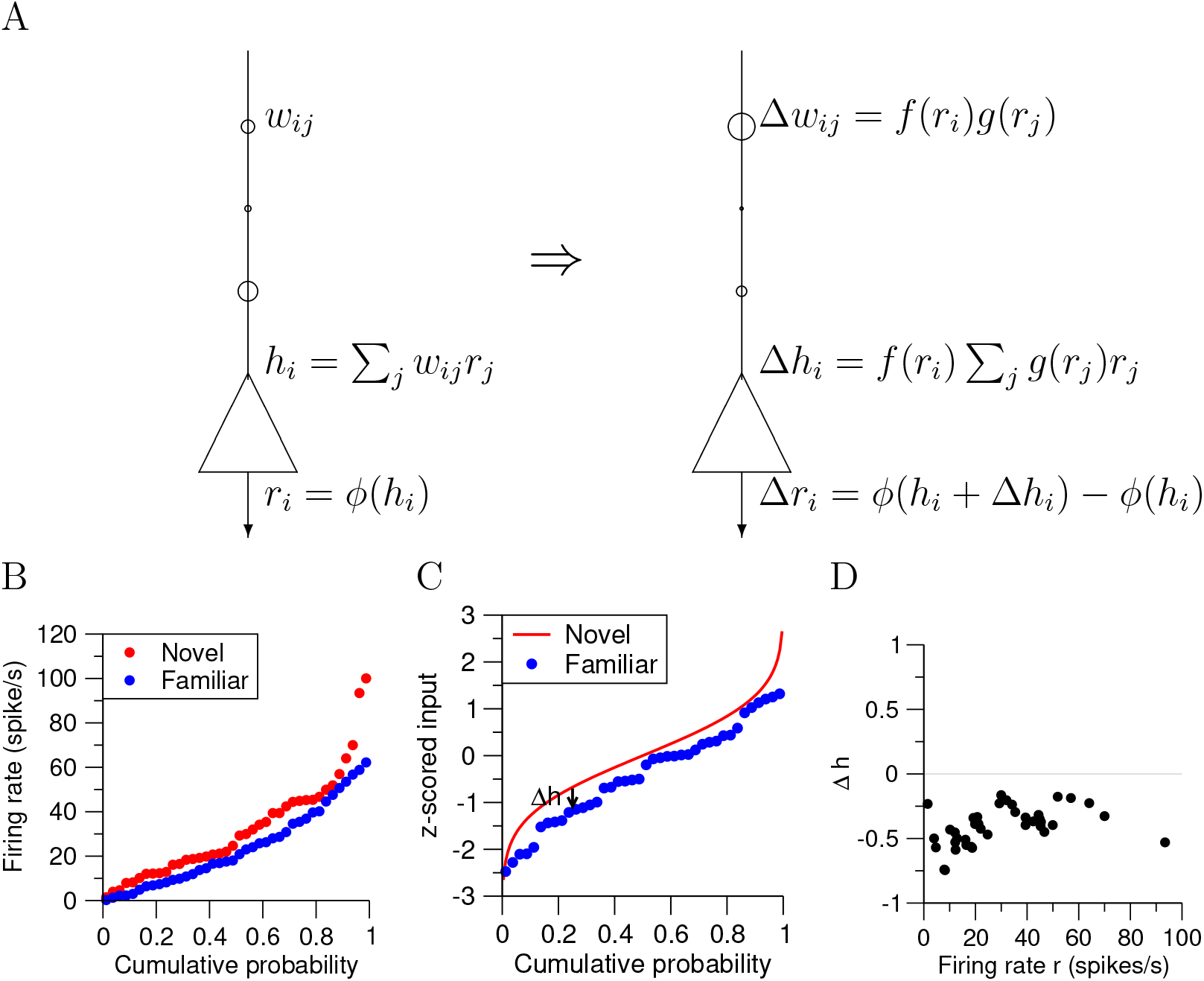
A. Sketch of hypothesized plasticity rule leading to changes of distributions of visual responses with familiarity. Synaptic efficacy changes Δ*w*_*ij*_ are given by a separable function of pre and post-synaptic rates *f* (*r*_*i*_)*g*(*r*_*j*_) The changes in total synaptic inputs *h*_*i*_(and consequently in visual response *r*_*i*_, which a function of total synaptic input *h*_*i*_ through the transfer function *ϕ*) with familiarity are proportional to *f* (*r*_*i*_), which describe the dependence of the plasticity rule on post-synaptic firing rate. B. Quantile functions of visual responses to novel and familiar stimuli of an IT neuron. C. Inferred quantile functions of total synaptic inputs for familiar stimuli, assuming distribution of inputs for novel stimuli is a Gaussian, for the same neuron shown in B. Changes in synaptic inputs Δ*h* are given by the difference between the two quantile functions. D. Changes in synaptic inputs as a function of visual response. In this particular neuron, changes are negative in the whole range of probed visual responses.

**Figure 6:**
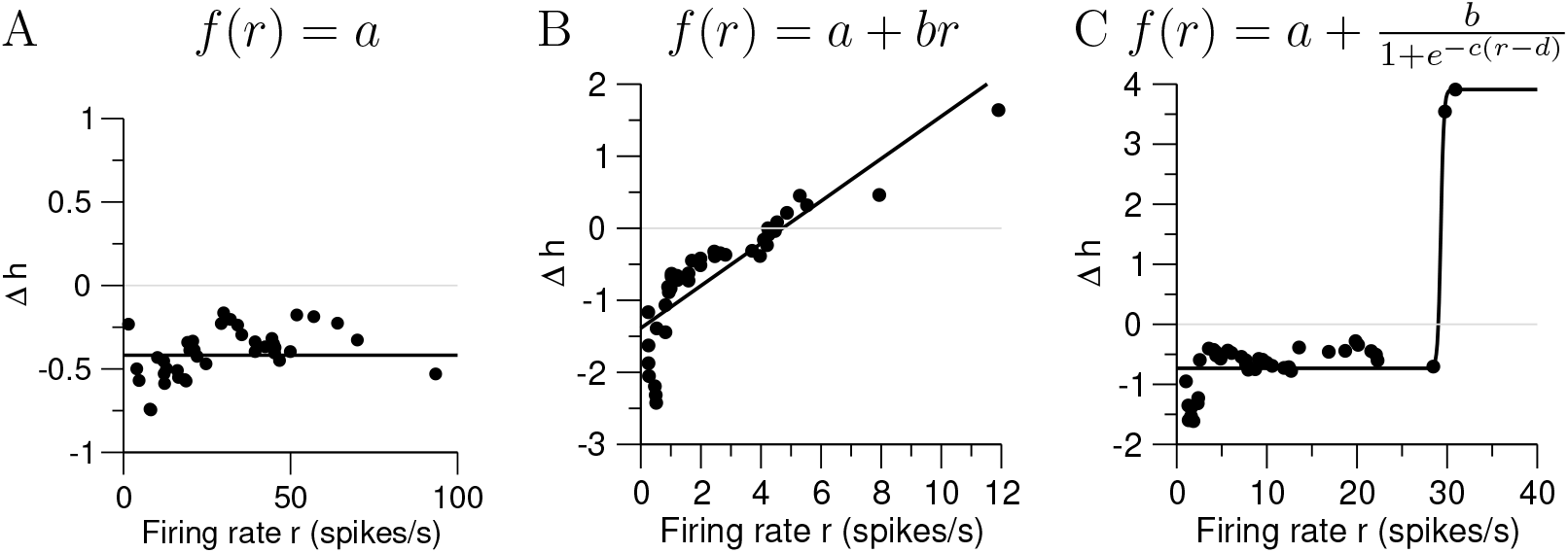
Fitting the dependence of synaptic plasticity on post-synaptic rate. A. Example neuron whose function Δ*h*(*r*) is well described by a constant, *f* (*r*) = *a*. It is the same neuron shown in Fig. 5. B. Example neuron whose function Δ*h*(*r*) is well described by a linear function, *f* (*r*)*a* + *br*. This neuron is consistent with a Hebbian rule with a linear dependence on post-synaptic firing rate. C. Example neuron whose function Δ*h*(*r*) is well described by a sigmoidal function, *f* (*r*)*a* + *b/*(1 + exp (− *c*(*r* − *d*))). This neuron is also consistent with a Hebbian rule, but with a step-function like dependence on post-synaptic firing rate.

Applying this procedure to the data, we found a strong heterogeneity of inferred plasticity rules. In some neurons, the inferred *f* (*r*) is approximately constant, with fluctuations around a value that can be positive or negative. In others, *f* (*r*) either decreases or increases with firing rate. To quantitatively characterize the dependence of *f* on post-synaptic firing rate, we fitted the inferred *f* (*r*) with three different types of functions: (i) a constant *f* (*r*) = *a*_*c*_; (ii) a linear function *f* (*r*) = *a*_*l*_ + *b*_*l*_*r*; and (iii) a sigmoidal function *f* (*r*) = *a*_*s*_ + *b*_*s*_*/*(1 + *exp* (− *c*_*s*_(*r* − *d*_*s*_))). Fig. (**??**) shows three examples of neurons, that are well described by a constant, linear, or sigmoidal fit, respectively.

To evaluate the significance of these inferred learning rules, we used a shuffling procedure. For each neuron, we generated 1000 samples by randomly shuffling visual responses for novel and familiar stimuli. We then fitted the shuffled data using the same functions as above, and generated a distribution of fitted parameters for the shuffles. We report in Tables 3, 2 the number of neurons whose fitted parameters are below the 5% or 1% percentiles of the shuffle distribution, and above the 95% and 99% percentiles of the shuffle distribution. For the linear fit, neurons whose fitted value of *a*_*c*_ is below/above the 5% percentile were termed Negative/Positive neurons, while neurons whose fitted value of *b*_*l*_ and/or *b*_*s*_ is below/above the 5% percentile were termed Anti-Hebbian/Hebbian neurons. Tables 3 and 2 show the numbers of neurons in each category, for visual responses computed in two different intervals an early interval (75-200ms after visual stimulus presentation) and the full visual response, respectively. We found in both cases that all these types of neurons were over-represented compared to reshuffled data, but to different extents. The largest category was the one of ‘Negative’ neurons, which in case of the full response is composed of 32% of all recorded neurons. The next largest category was composed of anti-Hebbian neurons, who were 19%/18% using linear/sigmoidal fits respectively. Other types of neurons were weakly represented - Positive neurons were only 6%, while Hebbian neurons were 8%/5% using linear/sigmoidal fits respectively. Our results reveal that learning rules inferred from the data are dominated by depression for all postsynaptic firing rates and this explains one of the neural correlates of familiarity - the gradual decrease in average firing rates observed in the data over long timescales.

**Table 2:**
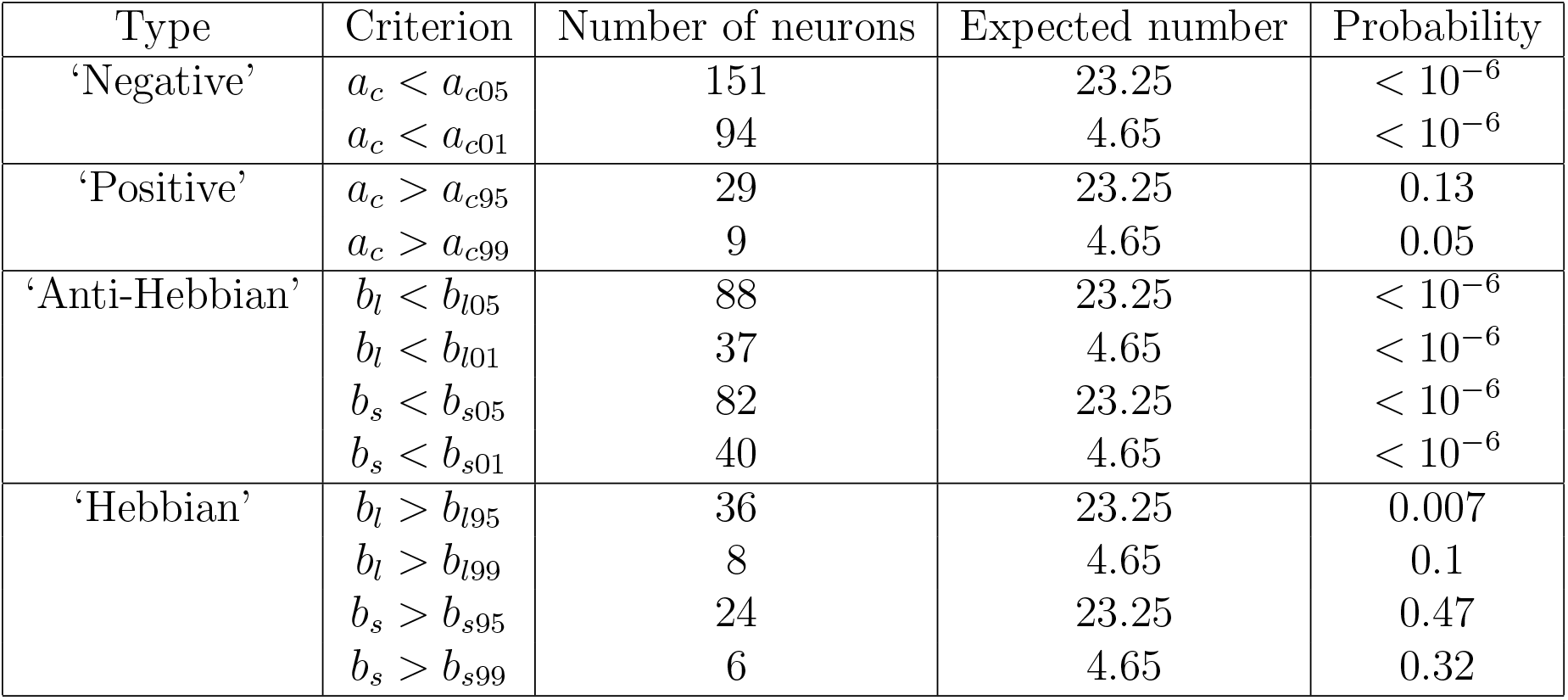
Number of neurons whose fitted parameters are significantly outside a 95% or 99% confidence interval, as determined by randomly shuffling visual responses. Visual responses computed from the full response. Note the much larger fraction of ‘negative’ neurons.

| Type | Criterion | Number of neurons | Expected number | Probability |
| --- | --- | --- | --- | --- |
| ‘Negative’ | $a_c < a_{c05}$ | 151 | 23.25 | $< 10^{-6}$ |
| | $a_c < a_{c01}$ | 94 | 4.65 | $< 10^{-6}$ |
| ‘Positive’ | $a_c > a_{c95}$ | 29 | 23.25 | 0.13 |
| | $a_c > a_{c99}$ | 9 | 4.65 | 0.05 |
| ‘Anti-Hebbian’ | $b_l < b_{l05}$ | 88 | 23.25 | $< 10^{-6}$ |
| | $b_l < b_{l01}$ | 37 | 4.65 | $< 10^{-6}$ |
| | $b_s < b_{s05}$ | 82 | 23.25 | $< 10^{-6}$ |
| | $b_s < b_{s01}$ | 40 | 4.65 | $< 10^{-6}$ |
| ‘Hebbian’ | $b_l > b_{l95}$ | 36 | 23.25 | 0.007 |
| | $b_l > b_{l99}$ | 8 | 4.65 | 0.1 |
| | $b_s > b_{s95}$ | 24 | 23.25 | 0.47 |
| | $b_s > b_{s99}$ | 6 | 4.65 | 0.32 |

**Table 3:** Coefficients of a quasi-poisson GLM model to fit neural firing rates: firing ∼ rate day number (1 to 8) + image category (N/IF) + neuron number.

| Predictor | Estimate | Std. Error | t-value | Pr(> \|t\|) |
| --- | --- | --- | --- | --- |
| day2 x NFIF | -0.03 | 0.03 | -0.89 | 0.37 |
| day3 x NFIF | -0.05 | 0.03 | -1.65 | 0.09 |
| **day4 x NFIF** | -0.07 | 0.03 | -2.32 | **0.01** |
| **day5 x NFIF** | -0.08 | 0.03 | -2.81 | **0.005** |
| **day6 x NFIF** | -0.11 | 0.02 | -4.33 | **$10^{-5}$** |
| **day7 x NFIF** | -0.09 | 0.02 | -3.78 | **$10^{-4}$** |
| **day8 x NFIF** | -0.06 | 0.02 | -2.68 | **$10^{-3}$** |

### 2.6 A recurrent network model with slow and fast plasticity time constants reproduces the multiscale dynamics of familiarity learning

Plasticity rules inferred from differences in statistics in neural responses to novel and highly familiar images reveal the asymptotic properties of the learning process in IT cortex but they do not contain any information about the timescales of learning. What types of plasticity rules can reproduce the dynamic evolution of familiarity observed in the data over long and short timescales? To investigate the time course of familiarization in the model, we employed a model in which each visual image presentation modifies the connectivity matrix using different variants of synaptic plasticity rules. The network model, together with synaptic plasticity rules, is described in detail in Methods. In this model, we derived equations that describes the average firing rate in response to a novel stimulus as a function of repetition number within a single recording session (see Methods). We started with a model that has a single synaptic plasticity time constant. This model could reproduce the time course of familiarization within a single session (full black curve in Fig. 7A) but for parameters constrained to data within a single session, was unable to reproduce the time course of familiarization across days (full black curve in Fig. 7B). This indicates that a single plasticity time constant does not effectively capture the dynamics of learning at both long and short timescales.

**Figure 7:**
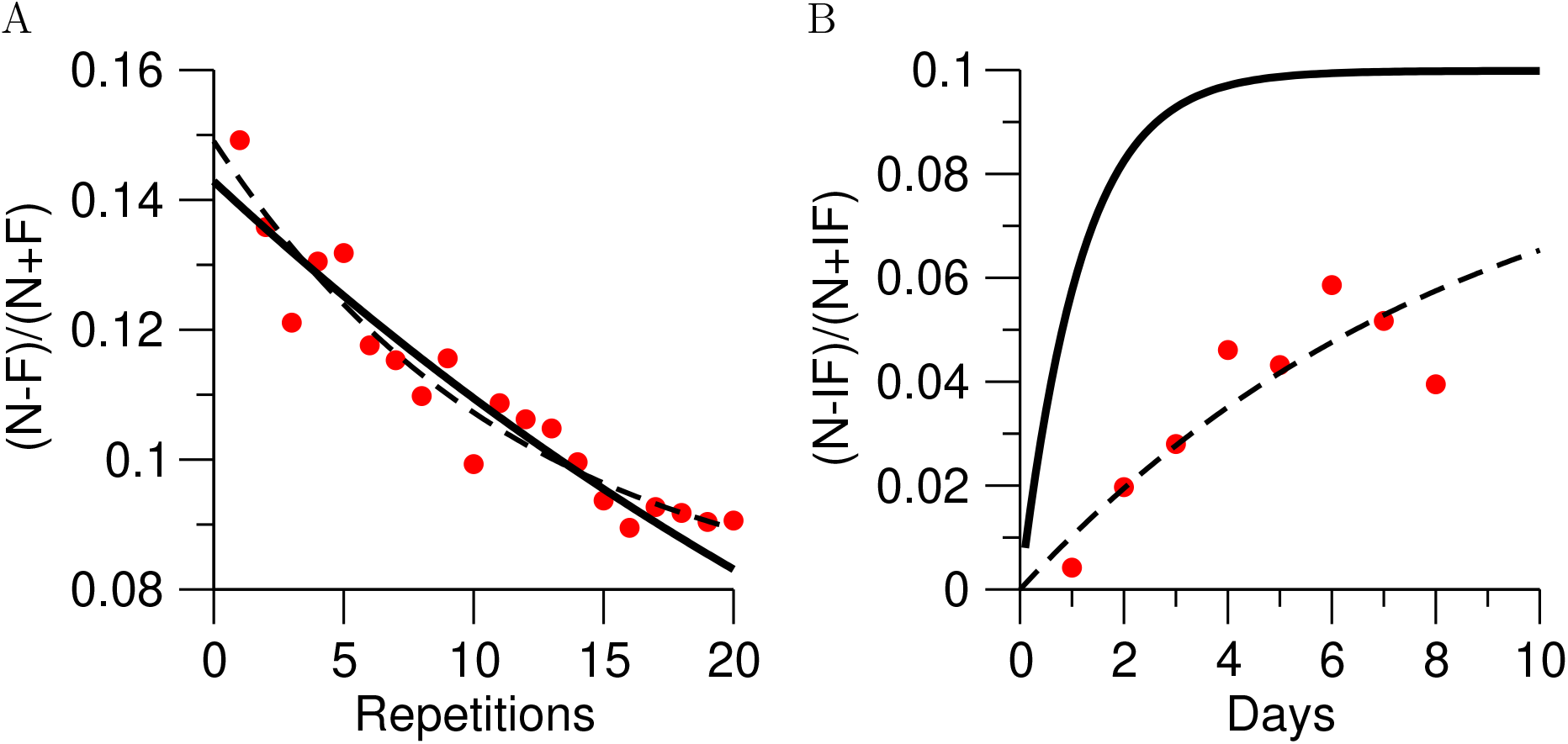
Time course of familiarization in a network model fitted to data. A. Average (*N* − *F*)*/*(*N* + *F*) index as a function of repetition number (Red circles: Data; Black full curve: Model with a single time constant; Dashed curve: Model with two time constants). B. Average (*N* − *IF*)*/*(*N* + *IF*) index as a function of number of previous days of exposure for intermediately familiar pictures. Red circles: Data; Black curve: Model with a single time constant, same parameters as in A. Dashed curve: Model with two time constants.

Our experimental results suggest that familiarity learning could occur on at least two different time scales. While average visually-evoked firing rates gradually decreased with familiarity over long timescales of days, spontaneous activity increased within single sessions over shorter timescales of hours. To test this idea, we then turned to a model that has two time constants: a fast time constant that modifies synaptic efficacies within single sessions; and a second slower time constant that describes a slower process of memory consolidation. With two time constants, the model could then capture both data within single sessions, and data across days (dashed curves in Fig. 7A-B). Thus, the neural data is consistent with a synaptic plasticity process that operates on two distinct timescales, one operating on the scale of minutes to hours (single sessions), the other on the scale of days.

## 3 Discussion

We demonstrate that the inferotemporal cortex maintains a detailed record of visual familiarity over timescales that span a 1000-fold range from tens to hundreds to thousands of image presentations. IT neurons encode familiarity through a progressive decrease in average firing rates, responding strongest to novel images, weaker responses to intermediately familiar images, and weakest to highly familiar images. While IT contained a robust code for familiarity at multiple timescales, the neural impacts of familiarity learning were slow to develop. Neural encoding of initially novel images became similar to highly familiar images gradually over a week (∼400 views) of repeated presentations, while representations of novel and familiar stimuli within single sessions remained quite distinct. Visually evoked firing rates to novel and familiar stimuli also decreased rapidly within sessions, accompanied by an increase in spontaneous activity. These differing patterns of neural activity at long- versus short-timescales suggest the hypothesis that visual learning is mediated by distinct mechanisms at different timescales. In support of this hypothesis, RNN models endowed with two synaptic plasticity rules, a slow time constant for longer timescales and a fast time constant for shorter timescales, replicated the observed neural dynamics of visual familiarity learning at multiple timescales.

Several studies have investigated the effects of visual experience in IT cortex and reported an average decrease in neural activity to repeated images and to a smaller extent, sharpening of tuning to visual features (Fahy et al., 1993; Fuster and Jervey, 1981; Woloszyn and Sheinberg, 2012a). According to this model, with repeated experience, neural populations encode a small number of preferred (maximally effective) familiar stimuli compared to unfamiliar stimuli. This sharpened tuning is believed to be implemented in the form of a winner-take-all network in which populations of neurons that are activated by stimuli show increased firing rates, whereas inactive or less active pools of neurons show decreased firing rates (Woloszyn and Sheinberg, 2012a). As a result, we would expect to see an increase in activity to the preferred stimuli and suppression of activity to all other stimuli. However, our results do not support this hypothesis since both mean and maximum firing rates (Figure S2) decrease over both long- and short-timescales, albeit at different rates.

A recent study found that the responses of excitatory neurons (but not inhibitory neurons) to their ‘best’ stimuli increased with long-term familiarity (Woloszyn and Sheinberg, 2012b). Moreover, Lim et al. (2015) found that this phenomenon could be captured quantitatively by a Hebbian learning rule in excitatory-to-excitatory connections, characterized by a non-linear dependence on the post-synaptic firing rate, that is dominated by depression. In contrast to these findings, the inferred plasticity rules were dominated by ‘negative’ learning rule (not Hebbian). To account for technical differences between the two studies, we matched distributions of firing rates and number of images used and still found that the results in the current study were consistent with negative learning rules. It’s possible that the two studies sampled from different parts of IT cortex or that history of training in animals could have resulted in different local circuits in IT cortex. In the current study, we used multi-electrode probes sampling in an unbiased way from dozens of neurons simultaneously whereas Woloszyn and Sheinberg (2012a) used single electrodes that were pre-screened to be visually selective and these differences could have also resulted in the discrepancy in inferred learning rules.

This study examines the dynamics and mechanisms of familiarity learning in IT and long and short timescales. We find that IT circuits support different mechanisms at different timescales with a fast plasticity time constant for short timescales (over hours) and a slow plasticity time constant for long timescales (over days). Our findings provide a data-driven approach to infer learning rules that account for the dynamics of cortical networks at both the short-term memory, but also the long-term memory timescales.

## 4 Methods

### 4.1 Experimental methods

#### 4.1.1 Subjects

Two male adult rhesus macaques (*Macaca mulatta*) weighing 12 kg and 14 kg were surgically implanted with a headpost and a recording chamber (20 mm inner diameter, Crist instruments). All surgeries followed sterile, aseptic technique with the animals under isoflurane anesthesia. All surgical and experimental procedures were in accordance with the University of Chicago’s Animal Care and Use Committee and US National Institutes of Health guidelines.

#### 4.1.2 Behavioral task and stimuli

Monkeys were trained to perform a RSVP dimming detection task in which they were required to detect and indicate (by releasing a manual touch bar) a subtle decrease in the luminance of the stimulus. The detection and report of image dimming encouraged monkeys to direct their attention to the presented stimuli. Each trial started when the animals grasped the manual lever and fixated at a central fixation spot for 500 ms. After gaze fixation was acquired, a series of images were presented at central fixation for 400 ms each with an inter-stimulus interval of 150 ms. On half of the trials, up to a maximum of 5 images were presented sequentially and dimming occurred on any of the five images with equal probability. After image dimming, monkeys were rewarded for releasing the touch-bar within 500 ms of dimming. On the remaining half of the trials, five images were always presented, and no dimming occurred. After image presentation, monkeys were rewarded for continuing to hold the touch-bar for 300 ms.

The stimuli used in this study were 150x150-pixel (4 degrees of visual angle) color images downloaded from Flickr using the program Bulkr (example images in Figure 1). The image set consisted of equal numbers of natural and man-made objects and scenes. Monkeys were first familiarized with a large library of 200 images that were viewed at least 1000 times during the dimming detection task. This image familiarization phase lasted for 3-5 months before neural recordings. After familiarization, in every recording session, 120 unique images that systematically varied in their familiarity were presented as monkeys performed the dimming detection task. Each recording session consisted of 40 novel, 40 intermediately familiar, and 40 highly familiar stimuli that were each presented at least 30 times within a single session. The novel images consisted of unique images that were never seen before a given recording session and were replaced with a new set of images for every subsequent recording session. The intermediately familiar images consisted of images that were viewed 50-500 times over a week during neural recordings and were retained over several consecutive recording sessions until they were cumulatively presented ∼500 times (∼8 days). The highly familiar images were randomly drawn from the previously familiarized library of images.

In the dimming detection task, monkeys were required to maintain fixation within ±2° of a 0.3° circular fixation point at the center of the screen for the duration of the trial. Eye movements were monitored and stored using an EyeLink infrared eye tracking system (SR Research Ltd., Ontario, Canada) at a sampling rate of 1 KHz. Precise control of task events, stimulus presentation, monitoring and storing of behavioral events and fluid-reward delivery were controlled via a MATLAB-based toolbox, MonkeyLogic. Stimuli were displayed using a 21-inch CRT monitor running at 75 Hz, 1280x1024 resolution, 24-bit color, positioned 57 cm from the monkeys’ eyes.

#### 4.1.3 Electrophysiological recordings

Structural magnetic resonance imaging scans (MRI) were used to identify area IT and to determine stereotaxic coordinates for recording chamber placement. IT recordings were targeted to the lower bank of the STS (areas TE, TEO) and the ventral surface of the inferior temporal cortex lateral to the AMT (area TE). IT recordings were conducted between AP coordinates 14-22 mm and ML coordinates 13-18 mm. Single units were recorded extracellularly with 24 channel V-Probes, a dura-piercing guide tube and were advanced using a NAN Microdrive system (NAN Instruments). Neurons were not pre-screened for visual selectivity. Neuronal waveforms were amplified, digitized, and stored for offline sorting to verify the quality and stability of neuronal isolation (Plexon Inc.).

#### 4.1.4 Data analysis and epoch-based analysis

All analyses were conducted on fixated images in all trials, excluding images with incorrect responses, fixation breaks, and early responses. All reported results are combined across two monkeys as results were qualitatively and quantitatively similar in both monkeys.

The baseline period was a 400-ms epoch prior to the onset of the first image on each trial. The image presentation period was a 400-ms epoch aligned to image onset, offset by 80 ms to account for neural response latency. Peri-stimulus time histogram plots for single neurons were averaged over all images in a given category (novel, familiar, intermediately familiar) and spike trains were convolved with a 20-ms boxcar kernel. For rank plots, firing rates were averaged over the image presentation period for each image, and images in each category were sorted by firing rates and plotted in ascending order. Mean peri-stimulus time histogram plots for neural populations were averaged across all visually-selective neurons and averaged over all images in a given category, and spike trains were convolved with a 20-ms boxcar kernel. Maximum peri-stimulus time histogram plots for neural populations were averaged across all visually-selective neurons with firing rates measured for the best image in each category (the image that elicited the highest firing rates), and spike trains were convolved with a 50-ms boxcar kernel (as opposed to the 20-ms smoothing window that was used for the mean PSTH plots since firing rates for a single image are inevitably noisier than the average). Mention how max firing rates are calculated? Mention SEM?

#### 4.1.5 Quantifying neuronal stability

The stability of recorded units was quantified by manually rating every recorded neuron with a score that ranged from 1-5. The stability score indicates the signal-to-noise ratio of a neuron, and depends on a number of factors such as the distance of the neuron from the electrode, length of the recording session, electrode drift, etc. A score of 5 indicates that the recorded unit was a single, well-isolated unit with little to no noise, and nearly all recorded spikes belonged to that unit. Conversely, a score of 1 indicates that the unit likely consisted of multiple single units that could not be resolved into independent units, or was significantly impacted by noise and electrode drift. For all analyses described here, we only included well-isolated units with stability score greater than or equal to three.

#### 4.1.6 Quantifying visual selectivity

Single neurons were defined to be visually selective if they responded with significantly different firing rates to at least one out of 120 images in the presented image set. Visual selectivity for each neuron was quantified using a linear model with image identities as categorical factors (i.e. 120 factors corresponding to 120 images), and was tested for significance with an F-test (*p <* 0.05). To account for global changes in firing rate due to familiarity that could interact with image selectivity, we used a linear model with image identity (1 to 120 images), repetition (1 to 50 repetitions), and image familiarity (novel, familiar, and intermediately familiar) as factors and we tested whether individual factors were significant using an F-test (*p <* 0.05). Tukey’s HSD test (*p <* 0.05) was used to further quantify whether neurons responded with significantly higher mean firing rates for novel, familiar, or intermediately familiar images.

### 4.2 Computational methods

In this section we describe the details of our network model, data analysis and mean-field analysis. They are extensions of models and methods introduced by Lim et al. (2015) and Pereira and Brunel (2018).

#### 4.2.1 Network model Assumptions

- External synaptic currents to neurons in ITC produced by novel images can be modelled by i.i.d. random Gaussian variables. The justification for this assumption is that neurons in cortex receive a large number of input synapses, so total synaptic inputs are likely to be well approximated by a Gaussian, by virtue of the central limit theorem.
- Synaptic currents that correspond to different images are statistically independent. This assumption is made for analytical tractability.
- Neural dynamics can be modelled by a ‘rate’ formalism. We focus on a rate model since the quantities of interest are (trial-averaged) firing rates of neurons in response to visual stimuli.
- Presentation of images lead to modifications of synaptic connectivity through an unsupervised firing rate based plasticity rule. The fact that synaptic strength can be modulated bidirectionnally by changes in firing rates of pre and post-synaptic neurons is well established in in vitro cortical preparations (see e.g. (Sjöström et al., 2001)), and such a rule has been shown to describe well changes in distributions of firing rates with familiarity (Lim et al., 2015).

##### Network model

We describe the dynamics of a network of *N* neurons in ITC by a system of rate equations (Dayan and Abbott, 2001; Hopfield, 1984), in which the firing rates of neurons in the network *r*_*i*_(*t*) (*i* = 1, …, *N*) evolve according to first order ordinary differential equations,

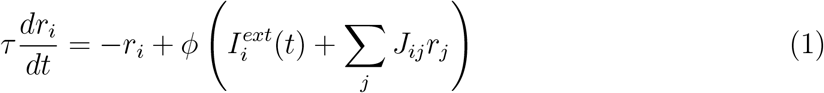

where *τ* is the time constant of rate dynamics (assumed to be on the order of 10ms (Reinhold et al., 2015)), *ϕ* is the f-I curve (transfer function) of neurons in the network, 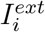 is the external input to neuron *i*, and *J*_*ij*_ is the recurrent connectivity matrix. Note that in the following we will only use the steady state solutions to these equations with a constant external input. Note also that in the following time will be measured in units of the interval between presentations of successive images. The justifications for using rate equations are that (1) the experimental data of interest are the mean firing rates of recorded neurons in ITC; (2) in many instances, the dynamics of networks of more realistic spiking neurons can be well approximated by such rate equations (Ostojic and Brunel, 2011; Sanzeni et al., 2020).

##### External inputs during presentations of images

External inputs during presentation of image *μ* are given by 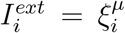 where 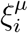 are i.i.d. random Gaussian variables, 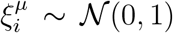. As in the experiment, images are separated in three categories: Familiar (*μ* ∈ ℱ) that have been shown many times in the past and continue to be presented every day; Novel (*μ* ∈ 𝒩) images that are presented during a single day; and Intermediately Familiar (*μ* ∈ ℐ) images that are presented during *n* successive days, where *n* varies from 2 to 8. The numbers of familiar/novel/intermediate familiar images presented on a given day are denoted by *p*_*F*_, *p*_*N*_ and *p*_*I*_, respectively, with *p* = *p*_*F*_ + *p*_*N*_ + *p*_*I*_.

##### Synaptic plasticity rule with a single time constant

We assume that an image presented during a trial *t* during a particular day *d* induces changes in synaptic connectivity, as

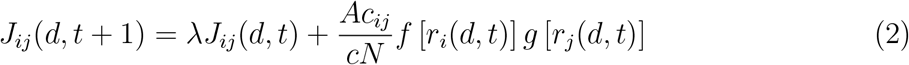

where 0 *< λ <* 1 is a decay parameter that determines the forgetting rate; *c*_*ij*_ = 0, 1 is a random Erdos-Renyi ‘structural’ connectivity matrix that determines the connectivity of neurons in the network, *c*_*ij*_ = 1 with probability *c*, 0 otherwise; *r*_*i*_(*d, t*) and *r*_*j*_(*d, t*) are the firing rates (visual responses) of post and pre synaptic neurons reached during the presentation of the image shown at time *t* on day *d*; And *f, g* are functions describing the dependence of the plasticity rule on post- and pre-synaptic rates, respectively, to be constrained by data. Note that this plasticity rule is similar to previously studied ‘palimpsest’ schemes, see e.g. (Mézard et al., 1986; Pereira-Obilinovic et al., 2023). For simplicity, we assume that the rates *r*^*μ*^ used to compute synaptic changes during presentation of an image *μ* are entirely determined by external inputs, 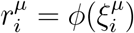 for all *i* when image *μ* is presented.

The function *g*(*r*) is required to have a zero average

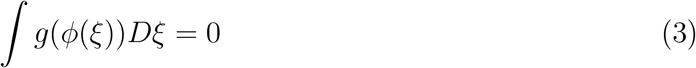

where *Dξ* is the Gaussian measure. This assumption is made so that the average synaptic changes are zero.

##### Imprinting of novel, intermediate familiar and familiar images in synaptic connectivity

With such a synaptic plasticity rule, the synaptic connectivity is at all times a linear superposition of traces left by all shown images,

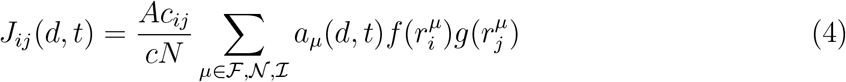

where *a*_*μ*_(*d, t*) denotes the strength with which a particular image *μ* has been imprinted in recurrent synaptic connectivity. *a*_*μ*_(*d, t*) depends on the history of presentations of the corresponding image *μ*. During an experimental session, each image is presented on a given trial with probability 1*/p*. Thus, for images that are presented on a given day, we can compute how the average imprinting strength ⟨*a*_*μ*_(*d, t*)⟩ evolves in time,

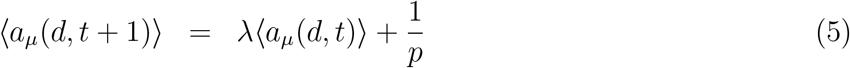

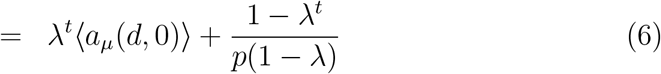

For an experimental session in which *T* images are presented in total, we have at the end of the experimental session

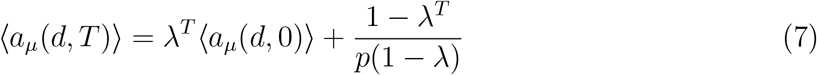

Using these equations, we can compute how the average imprinting strengths evolve in time during sessions, for the three classes of images:

- For novel images, *μ* ∈ 𝒩, the initial imprinting strengths are ⟨*a*_*μ*_(*d*, 0)⟩ = 0, and

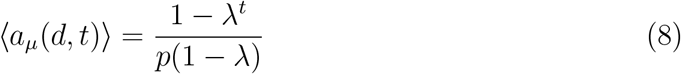

- For intermediate familiar images *μ* ∈ ℐ, the initial imprinting strengths can be obtained by recursion using Eq. (7),

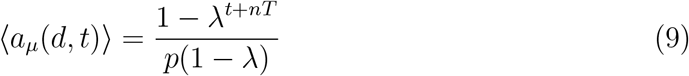

where *n* is the number of previous days of exposure with such images.

- Finally, the average imprinting strengths for familiar images are obtained using the *n* → ∞ limit in the above equation, i.e.,

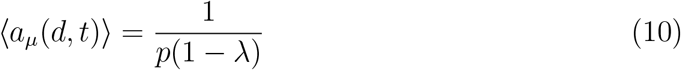

In the following we will also need the average these imprinting strengths over sessions, denoted *A*_0_, *A*_*n*_, and *A*_*∞*_ for novel, intermediate familiar with *n* days of previous exposure and familiar, respectively. These averages are obtained from taking the average of *λ*^*t*^ from *t* = 0 to *t* = *T* − 1, i.e. by replacing *λ*^*t*^ with (1 − *λ*^*T*^)*/*(*T* (1 − *λ*) in Eqs. (8,9,10) respectively.

##### Average visual responses as a function of time within a session, and familiarity

We now turn to how average visual responses depend the history of presentations of a particular image. With network dynamics given by Eq. (1) and a synaptic connectivity matrix given by Eq. (4), the steady state rate of neuron *i* during presentation of image *μ* on day *d* at time *t* is

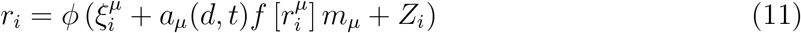

where the first term in the argument of the transfer function *ϕ* is the external input, the second term (proportional to *a*_*μ*_) is proportional to the embedding strength of the shown image in the connectivity matrix and the overlap *m*_*μ*_, defined as

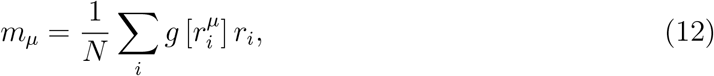

and the last term is a noise term due to other patterns stored in the connectivity matrix. From Eqs. (11,12), we obtain in the 1 ≪ *cN* ≪ *N* limits mean-field equations, the mean rate *R*_*μ*_(*d, t*), and the overlap:

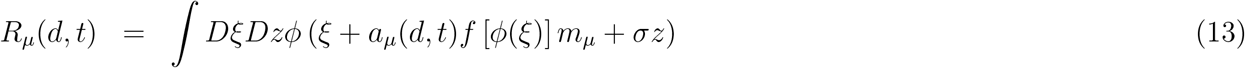

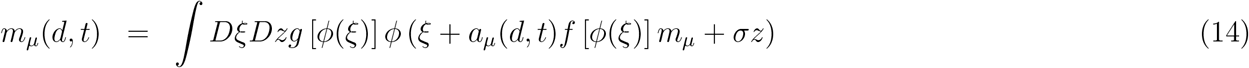

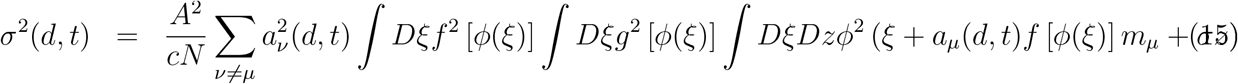

Assuming that the embedding strengths have a small impact on the average rate, we can Taylor expand the above equations in *a*_*μ*_ and obtain up to first order in *a*_*μ*_

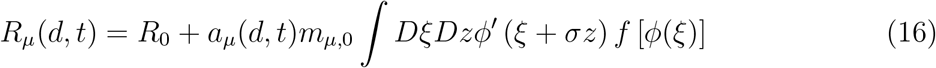

where *R*_0_ is the average firing rate when a novel stimulus is shown,

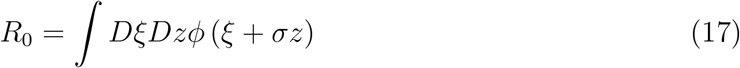

and *m*_*μ*,0_ is given by

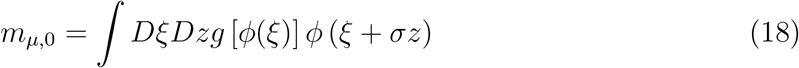

##### Synaptic plasticity rule with two timescales

The observation of two different timescales of effects of familiarity on mean firing rates, and the inability of the simplest one time scale model to fit the data, motivates the introduction of two timescales. We assume that the imprinting strengths associated with each image *a*_*μ*_ obey two different dynamics. Each image has a probability *ρ* of being consolidated in memory - if it is consolidated, the corresponding imprinting strength is *a*_*c*_. This probability is *ρ* = 0 for novel images; 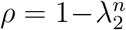 for intermediately familiar images, with *n* days of previous exposure to the corresponding image; and *ρ* = 1 for familiar images. Thus, the parameter *λ*_2_ describes the time scale of consolidation. Images that are not consolidated follow the same dynamics as Eq. (2), replacing *λ* by *λ*_1_. We also assume for simplicity that these changes are washed away in between sessions - thus, either images are consolidated the next day, or there is no memory of them in the synaptic connectivity. This leads to the following average imprinting strengths for the three types of images:

- For novel images, *μ* ∈ 𝒩,

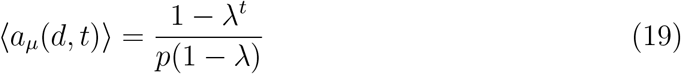

- For intermediate familiar images *μ* ∈ ℐ,,

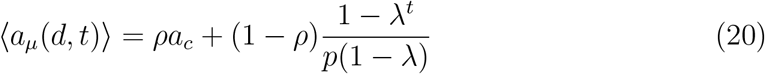

where *n* is the number of previous days of exposure with such images.

- Finally, the average imprinting strengths for familiar images are

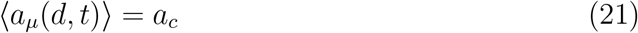

#### 4.2.2 Obtaining *f* and *λ* from the data

To obtain *f* in individual neurons, we follow the analysis introduced by Lim et al. (2015) to compare the statistics of the total synaptic currents in the presence of novel and familiar stimuli. From Eq. (11), we see that the average synaptic input to neuron *i* in response to a novel stimulus *μ* ∈ 𝒩, conditioned by this stimulus is simply 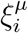, while for a familiar stimulus *μ* ∈ ℱ it is 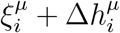 where

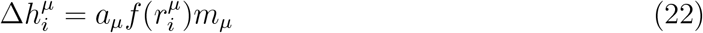

Eq. (22) shows that we can infer the dependence of the learning rule on the post-synaptic firing rate *f* (*r*) from the dependence of Δ*h* on firing rate *r*.

To infer the transfer function *ϕ* and the dependence of the post-synaptic plasticity rule on firing rate *f* of a particular neuron *i* in the data set, we perform the following steps (Lim et al., 2015):

- Compute the quantile function of visual responses to novel images 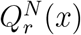, by rank ordering the visual responses, 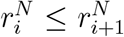, for *i* = 1, …, *n*_*N*_, and setting 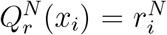, where *x*_*i*_ = (2*i* + 1)*/*(2*n*_*N*_). We can derive similarly the quantile function of visual responses to familiar images, 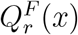.
- The transfer function *ϕ* is then obtained thanks to the relationship 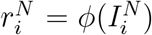, and our assumption that input currents when a novel image is presented, are drawn from a Gaussian distribution with zero mean and unit variance. Thus, we can use the quantile function (inverse c.d.f) of a standard Gaussian 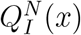, and find for each quantile *x*_*i*_, 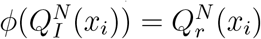.
- Given the transfer function *ϕ*, and the quantile function of visual responses to familiar images 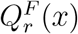, we can now obtain the quantile function of inputs for familiar images, 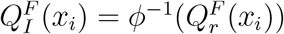. To compute the inverse of the transfer function *ϕ*, we need to interpolate (and extrapolate) the function *ϕ* which up to this point has been computed only on a discrete set of *n*_*N*_ quantiles. For a given firing rate 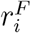 in response to a familiar stimulus, there are three possible scenarios:

1. If 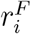 is non-zero, and within the range of novel firing rates 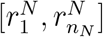, we interpolate linearly the transfer function between the two quantiles *k, k* + 1 that are such that 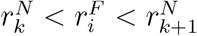;
2. If 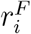 is non-zero and outside the range of novel firing rates, we need to extrapolate the transfer function outside of the range 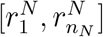. When 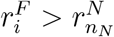, we pick the nearest quantile *i* such that 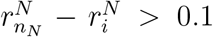, and use these two quantiles to extrapolate linearly *ϕ* above 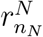. Likewise, when 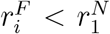, we pick the nearest quantile *i* such that 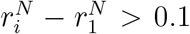, and use these two quantiles to extrapolate linearly *ϕ* below 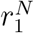.
3. If 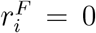, and there at least two stimuli for which novel responses were zero, then there is some ambiguity as to which current we should use for this quantile. A natural choice is to use the quantile which is the closest to *i* - in particular if 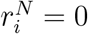, then we choose 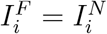.

- We now identify Δ*h*, for each quantile *x*_*i*_, as the difference between the two quantile functions 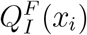 and 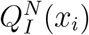. Finally, we can compute Δ*h* as a function of rate, as

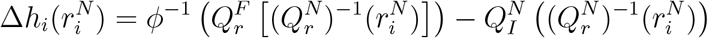

This procedure is described graphically in Fig. 5.

We fitted Δ*h*_*i*_(*r*) obtained as explained above using the following simple functions

- Constant

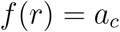

- Linear

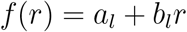

- Sigmoidal

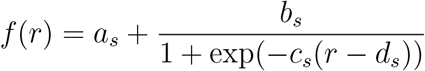

Note that both linear and sigmoidal fits with *b*_*l*_(*b*_*s*_) *>* 0 are consistent with Hebbian plasticity, while *b*_*l*_(*b*_*s*_) *<* 0 is consistent with anti-Hebbian plasticity. Note also that sigmoidal curves were used by Pereira and Brunel (2018) to fit data from Woloszyn and Sheinberg (2012b).

## 5 Supplementary Information

**Supplementary figure S1.**
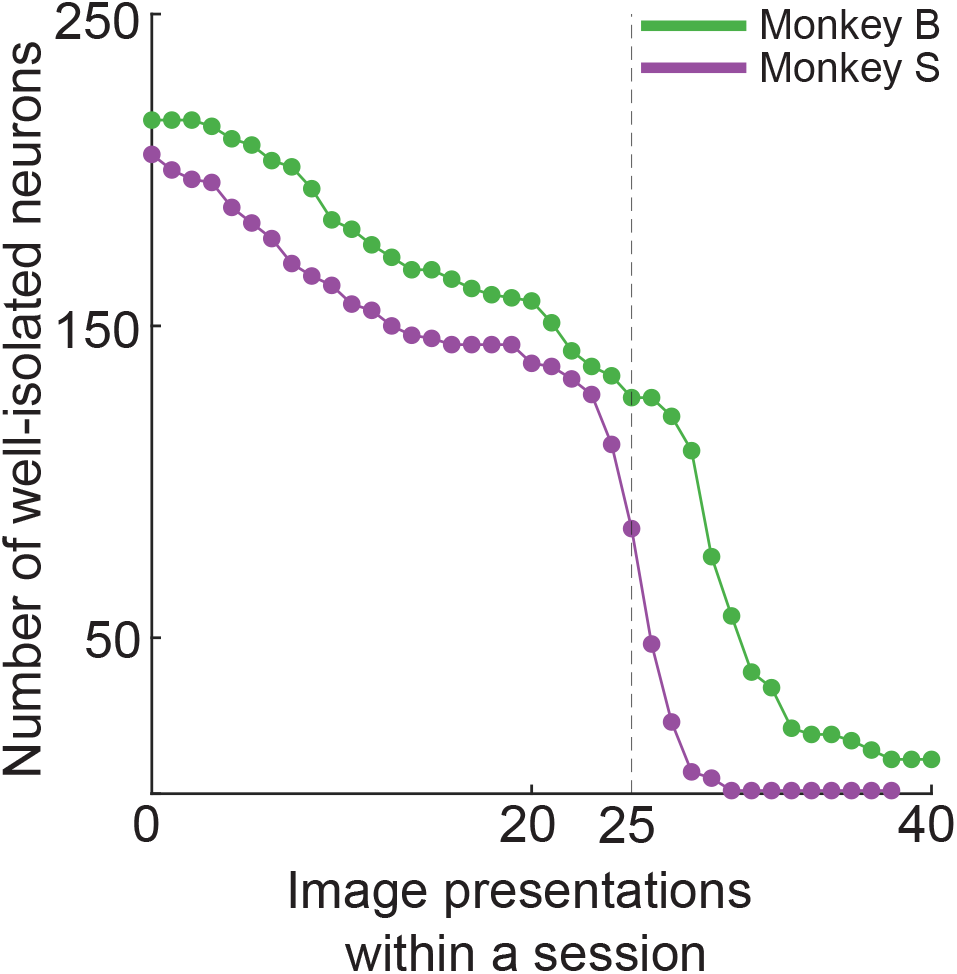
Cumulative neural yield within recording sessions. Plot of number of well-isolated neurons across image presentations within recording sessions. The neural yield drops steeply at about 25 image presentations within a session. Dashed line indicates the criterion for including neurons that were well-isolated from the start of a session up to 25 image presentations of all novel and highly familiar images within the same recording session.

**Supplementary figure S2.**
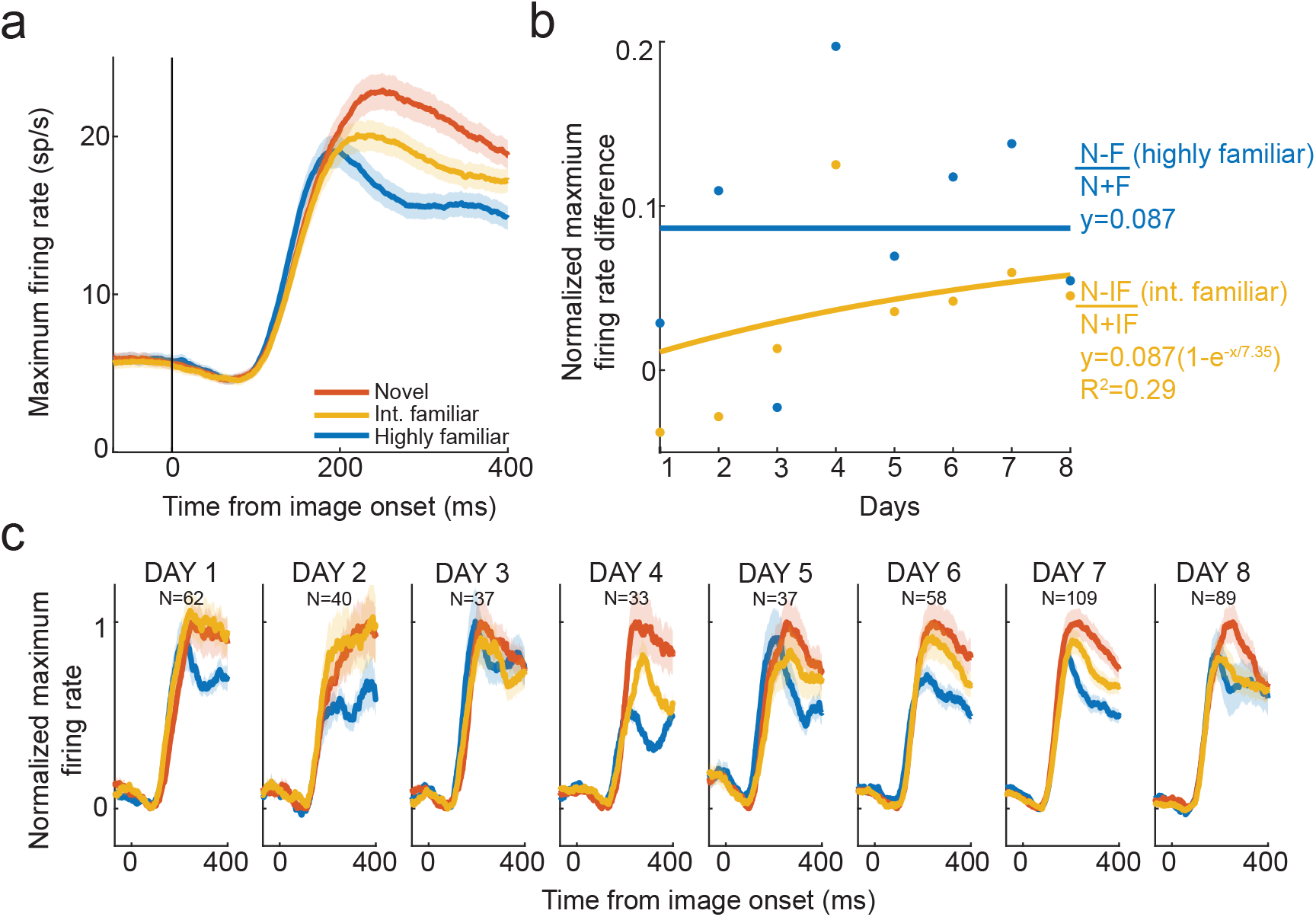
Long-term image familiarity evolves over multiple days in IT neural populations. (a) Peri-stimulus time histogram (PSTH) of visually selective IT neurons (N=465) to the best novel (seen 10s of times), best intermediately familiar (seen 100s of times), and best highly familiar images (seen 1000s of times). Maximum firing rates are strongest for novel, followed by intermediately familiar, and weakest for highly familiar images. (b) Normalized firing rate difference between responses to the best novel and best intermediately familiar images ((N-IF/N+IF) in orange) as a function of number of days of viewing intermediately familiar images. The blue curve represents the normalized firing rate difference between responses to the best novel and the best highly familiar images (N-F/N+F) as a function of number of days of viewing highly familiar images. Positive values indicate that neural responses to novel images are greater than neural responses to both intermediately familiar and highly familiar images. Over multiple days of viewing, the normalized difference between novel and intermediately familiar images increases, following an exponential function with an asymptote that reaches the constant value of highly familiar images. (c) Normalized PSTHs to novel, intermediately familiar and highly familiar images on each day. The difference between novel and intermediately familiar images increases over days, reaching statistical significance first on day three and staying significant until day eight (*p <* 0.05, GLM). For each day, the curves represent the PSTH pooled across all neurons recorded on that day, and over all images in a given category. Firing rates on each day were normalized by subtracting the baseline at t=0 and dividing by peak minus baseline, with both peak and baseline measured on mean activity combined across all images in a category. In both (a) and (c), spike trains were convolved with a 50-ms causal boxcar kernel.

